# Using hypertemporal Sentinel-1 data to predict forest growing stock volume

**DOI:** 10.1101/2021.09.02.458789

**Authors:** Shaojia Ge, Erkki Tomppo, Yrjö Rauste, Ronald E. McRoberts, Jaan Praks, Hong Gu, Weimin Su, Oleg Antropov

**Affiliations:** Nanjing University of Science and Technology, School of Electronic and Optical Engineering, Department of Electronic Engineering, 210094 Nanjing, China; Department of Forest Sciences, University of Helsinki, Latokartanonkaari 7, P.O. Box 27 FI-00014 Helsinki, Finland; Aalto University, Department of Electronics and Nanoengineering, P.O. Box 11000, FI-00076 AALTO, 02150Espoo, Finland; VTT Technical Research Centre of Finland, P. O. Box 1000, FI-00076 VTT, Espoo, Finland; University of Minnesota, Department of Forest Resources, Saint Paul, MN 55108, USA

**Keywords:** synthetic aperture radar, growing stock volume, boreal forests, Sentinel-1, support vector regression, random forests regression

## Abstract

In this study, we assess the potential of long time series of Sentinel-1 SAR data to predict forest growing stock volume and evaluate the temporal dynamics of the predictions. The boreal coniferous forests study site is located near the Hyytiälä forest station in central Finland and covers an area of 2,500 km^2^ with nearly 17,000 stands. We considered several prediction approaches (linear, support vector and random forests regression) and fine-tuned them to predict growing stock volume in several evaluation scenarios. The analyses used 96 Sentinel-1 images acquired over three years. Different approaches for aggregating SAR images and choosing feature (predictor) variables were evaluated. Our results demonstrate considerable decrease in RMSEs of growing stock volume as the number of images increases. While prediction accuracy using individual Sentinel-1 images varied from 85 to 91 m^3^/ha RMSE (relative RMSE 50-53%), RMSE with combined images decreased to 75.6 m^3^/ha (relative RMSE 44%). Feature extraction and dimension reduction techniques facilitated achieving the near-optimal prediction accuracy using only 8-10 images. When using assemblages of eight consecutive images, the GSV was predicted with the greatest accuracy when initial acquisitions started between September and January.

**Highlights:**

- Time series of 96 Sentinel-1 images is analysed over study area with 17,762 forest stands.
- Rigorous evaluation of tools for SAR feature selection and GSV prediction.
- Improved periodic seasonality using assemblages of consecutive Sentinel-1 images.
- Analysis of combining images acquired in “frozen” and “dry summer” conditions.
- Competitive estimates using calculation of prediction errors with stand-area weighting.

## 1. Introduction

Space-borne synthetic aperture radar (SAR) is a versatile Earth Observation tool capable of providing fine resolution, terrain-related data regardless of weather conditions. The data are particularly useful in areas with persistent cloud cover and that are close to polar regions. SAR also has a long history of use for forest remote sensing where models of relationships between forest structural parameters and measured backscatter signatures are essential (Sinha et al., 2015). SAR has been shown to be sensitive to forest structure variables including, and most importantly, above-ground tree biomass (AGB) (t/ha) and growing stock volume (GSV) (m^3^/ha) (GFOI, 2014). Depending on the sensor wavelengths, the relationship can saturate quickly and can additionally be affected by environmental and ionospheric conditions. Once signal saturation is reached, the radiometric data are no longer useful for volume or biomass prediction beyond the “saturation point”, particularly when only one image is used. One result is that data from the commonly used C- and X-band space-borne SAR systems are widely considered suboptimal for large area forest inventory and mapping (GFOI, 2014).

Cross-polarised (cross-pol) backscatter demonstrates greater sensitivity to forest biomass than co-polarised (co-pol) backscatter, although multiple polarizations are recommended for use in forest biomass mapping algorithms. L-band SAR is useful for discriminating among regrowth stages and estimating biomass in forests with small to moderate biomass levels (40-150 t/ha). Dual polarization and dual-season coverage are required. C-band SAR is mostly useful for forests with very small biomass levels (30-50 t/ha) when a single SAR intensity image is used. The shorter wavelengths have higher extinction and thus limited penetration within the forest layer (compared to L- and P-bands), although multi-temporal and texture analysis of fine resolution C-band data may provide useful inputs (Sarker et al., 2013; Bourgoin et al., 2018; Chen et al., 2019; Reis et al., 2019; Tomppo et al., 2019).

On the other hand, C-band radar data acquired by multiple space-borne sensors including historically ERS-1, ERS-2 and Envisat ASAR, presently RADARSAT-2 and ESA Sentinel-1, and since recently RADARSAT Constellation represent attractive sources of information for timely assessment of forest cover due to global coverage (in case of ASAR and Sentinel-1) and frequent revisits. This is particularly important for Sentinel-1 satellites (Torres et al., 2012) with data offered free of charge to users.

To date, multitemporal approaches for GSV estimation using L-band SAR imagery in boreal forests are well-demonstrated, particularly with JERS (Rauste, 2005; Kurvonen et al., 1999; Santoro et al., 2006), and ALOS PALSAR (Cartus et al., 2012; Antropov et al., 2013; Antropov et al., 2017). Methodological studies using long time series of C-band SAR intensity data for GSV mapping over boreal forests have been relatively rare (Kurvonen et al., 1999; Santoro et al., 2011; Tomppo et al.,2019; Pulliainen et al., 1999) and while the common understanding is that C-band data are suboptimal compared to L- and P-band data (GFOI, 2014), however due to availability of C-band data (Envisat ASAR in Global Monitoring mode), wide-area maps of GSV were produced in boreal forests (Santoro et al., 2011). Historical context of these studies is given further in Section 1.1.

Some research has been conducted on the fusion of Sentinel-1 data with SAR data from instruments operating at different wavelength as well as optical data. Particularly, the combination of Sentinel-1, ALOS-2 and Sentinel-2 data produced greater accuracies, although the Sentinel-1 role was relatively limited (Laurin et al., 2018). Another study compared single-date and multi-temporal Sentinel-1 data and demonstrated that multi-temporal Sentinel-1 substantially improved AGB estimation, but still with a much smaller adjusted R^2^ because of the limitations of the short wavelengths (Huang et al., 2018). When using the Water Cloud model (Attema & Ulaby, 1978) based approach for AGB prediction from ASAR hypertemporal data, relative root mean square errors (rRMSEs) in the range 34.2 – 48.1% were achieved for for 1 km pixel predictions (Santoro et al., 2011). Note that rRMSE is defined as the percentage ratio of root mean square error (RMSE) and the mean of the response variable *y* in the validation data set. GSV predictions were more accurate when averaging over neighbouring pixels. Use of multiple stand-level features was instrumental to achieve an rRMSE of 30% for a Finnish test site (Tomppo et al., 2019). There, stand-level backscatter intensity and other stand-level features, such as standard deviation of the intensities and the averages of the ratios of the intensities of the images from different dates, were used as predictor variables to produce large GSV prediction accuracies. Several sets of RADARSAT-2 images were studied using machine learning approaches in combination with other SAR and optical data using various features and machine learning approaches achieving rRMSE of 42-47% (Stelmaszczuk-Górska et al., 2018).

The key question when using remotely sensed data for terrain related variable predictions are 1) how quickly and how often the data are available, for instance, for damage monitoring, 2) what is the best season for image acquisition with respect to seasonal variation of the vegetation when the criterion is the accuracy of the predictions, 3) how many images are needed to obtain the maximum accuracy with a particular type of images, and 4) what feature selection approaches can be used to select the most useful predictor variables to maximise GSV prediction accuracy.

### 1.1. Background of multitemporal C-band SAR studies for GSV prediction

Parametric semiempirical and WCM based models have been used with historical C-band SAR data, such as ERS-1 and Envisat ASAR, to predict forest stem volume and GSV (Pulliainen et al., 1996; Pulliainen et al., 1999; Santoro et al., 2011). Historical detailed studies of temporal dynamics of C-band SAR backscatter in boreal forests are scarce, whereas in notable examples (Pulliainen et al., 1996; Pulliainen et al., 1999)) the SAR data were represented by ERS-1 single-pol VV imagery at 23 nominal incidence angle. In Pulliainen et al. (1996), the auxiliary parameters of a semiempirical model were estimated separately from each processed SAR image, and then a vector of backscatter measurements for each areal-unit was iteratively inverted to predict forest stem volume. Correlation analysis with reference plot-level stem volume was used to select suitable images. In Pulliainen et al. (1999), parameters of a semiempirical model and stem volume were estimated simultaneously for each SAR image. Separate stem volume estimates were later combined using a linear regression equation determined using the training data set. Also, use of multiple linear regression for optimally combining predictions from individual SAR images was reported also in (Kurvonen et al., 1999).

Overall during the last two decades, a popular approach in forest biomass mapping is represented by initially predicting GSV using WCM for individual SAR images and further combining those predictions via optimal weighting of the predictions (Santoro et al., 2011). Weights can be derived using image-wise accuracy statistics in the presence of reference data, or auxiliary datasets (Santoro et al., 2011; Santoro & Cartus, 2018).

More recently, nonparametric approaches (random forests, support vector regression) using features calculated from multiple C-band SAR images were tested, often in combination with L-band and optical images (Laurin et al., 2018). Using features calculated from several SAR images as independent variables in GSV prediction has rarely been studied ((Stelmaszczuk-Górska et al., 2018; Tomppo et al., 2019) and typically included not only SAR intensity but also other textural parameters. Temporal dynamics of GSV predictions based on long time series of consecutive C-band SAR data were not investigated, and only few feature selection approaches were used (such as random forests based ranking, and simple correlation to forest biomass).

Availability of long time series of freely available Sentinel-1 dual-pol (VV, VH) data enables more in-depth exploration of different ways of multitemporal SAR data processing in the context of GSV prediction. Studying various ways of combining C-band images via feature selection to predict GSV, as well as studying temporal dynamics of GSV predicted using selected features (including analysis of long sets of consecutive SAR images) has become possible.

While combining Sentinel-1 SAR imagery with other remotely sensed data is common, the effect of seasonal dynamics of Sentinel-1 SAR dual-pol data on GSV predictions in boreal forests has not yet been reported. Availability of very long time series, presence of two polarizations (co- and cross-polarizations) and access to better reference data motivates more in-depth exploration of seasonal dynamics and its influence on GSV predictions. The only report mentioning Sentinel-1 time series seasonality has been (Dostálová et al., 2018) where a forest/nonforest classification was attempted. However, the authors did not elaborate on the temporal dynamics, the length of the time series or its influence on classification accuracy as pixel-wise intensities of whole SAR image stack were used as a feature vector in classification.

A general conclusion is that the use of a single Sentinel-1 image does not lead to accurate forest biomass mapping. Therefore, the present research concentrates on the fusion of Sentinel-1 and L-band SAR or optical satellite data, as well as investigating the potential of multitemporal Sentinel-1 datasets. The performance of multitemporal Sentinel-1 approaches is not yet satisfactory, especially at stand-level, although aggregation of estimates to kilometre or county level is more accurate. Most critically, in-depth analysis of the potential of very long time series of Sentinel-1 data and their seasonal fluctuations (dynamics) and the effect of the fluctuation on the accuracy of GSV prediction is lacking, although the data for multiple years are readily available. Furthermore, a statistically sound method for optimising image and season selection for purposes of increasing prediction accuracy using dual-pol Sentinel-1 data has not been demonstrated over boreal forests. Reported studies of feature selection are relatively scarce, disparate and not exhaustive. This motivates us to address these knowledge gaps. In this study, we use representative reference forest stand-level data to study various GSV prediction models and feature selection approaches, including those not studied previously with SAR data.

### 1.2. Study goals and the contributions of the study

In this study we expand on a lineage of forest biomass mapping using multitemporal radar imagery by evaluating the use of large numbers of Sentinel-1 images for a lengthy time period for Finnish boreal forests. We focus on one key variable, GSV (m^3^/ha), defined here as the above-ground volume of living stems above stump to the stem top over a specified area. Included are the stem volumes of living trees, standing or lying with a diameter at breast height (dbh) of more than 0 cm and height of more than 1.30 m where dbh is measured. Note that dbh is measured at the height of 1.3 m. GSV is strongly correlated with other forest variables, such as tree basal area, AGB and total tree carbon content. The latter is approximately (0.3–0.4 t/m^3^)×GSV in boreal forests, depending on the tree species (Karjalainen & Kellomäki (1996), Tomppo (2000)). Thus, our results can be compared to the results from other relevant studies that use SAR to predict tree biomass for boreal forests.

Four specific questions are addressed in this study:

1. what is the seasonal variation of RMSE of stand-level GSV predictions?
2. what is the value of adding multitemporal imagery compared to single images?
3. how many images are necessary for achieving optimal accuracies and in exactly what manner?
4. what is the optimal season for Sentinel-1 image acquisition to maximise GSV prediction accuracy also when including sets of consecutive images?
5. which feature selection and GSV prediction methods give the greatest GSV prediction accuracies.

The primary and overarching objective of the study was to develop a method for using Sentinel-1 imagery to maximize GSV prediction accuracy. Three supporting and subordinate objectives were addressed: (1) to analyse the effect of the acquisition date, weather conditions and number of Sentinel-1 images on the GSV prediction accuracy, (2) to test and develop feature selection and transformation methods for backscattering coefficients and multitemporal features, such as, principal component analysis, radiometric contrast, mutual contrast, Lasso and genetic algorithm based optimization of the feature weights, and the effects of the features and transformations on the prediction accuracies, and (3) to test a few commonly used optional forest parameter prediction methods and their performances in GSV prediction.

These objectives were addressed using extensive analyses of a 3.5-year stack of Sentinel-1 images acquired over a large test site representative of boreal forestland. Both established and advanced statistical and machine learning approaches were used for feature selection and forest biomass prediction, particularly k-NN using genetic algorithm based feature optimization improved k-NN (ik-NN) (3.2.5), random forests (RF) and support vector regression (SVR). With management inventory needs in mind, we focused on stand-level GSV predictions and their error estimates, not necessarily on large area level estimates and their error estimates. The criterion was mean standard error at stand-level when using separate validation data. Note that estimators with large standard errors at stand-level may produce consistent large area estimators while precise stand-level estimators may be biased large area estimators.

## 2. Materials

### 2.1. Study site

The 50-km × 50-km study area in Southern Finland is centred around the Hyytiälä Forestry Field station of the University of Helsinki with centre coordinates: 61.8°N, 24.3 °E (Figure 1). The study site includes forest, agricultural land, lakes, and several population centres. The forests are dominated by Norway spruce (Picea abies (L) H. Karst.), Scots pine (Pinus sylvestris L.), and birch (Betula pendula Ehrh. and Betula pubescens Roth). The soils are mainly glacial drift, but sandy soils are also common as well as clay, especially in agricultural areas. Mires and open bogs are also common. The area is relatively hilly with terrain variation ranging between 95 and 230 meters above sea level.

**Figure 1:**
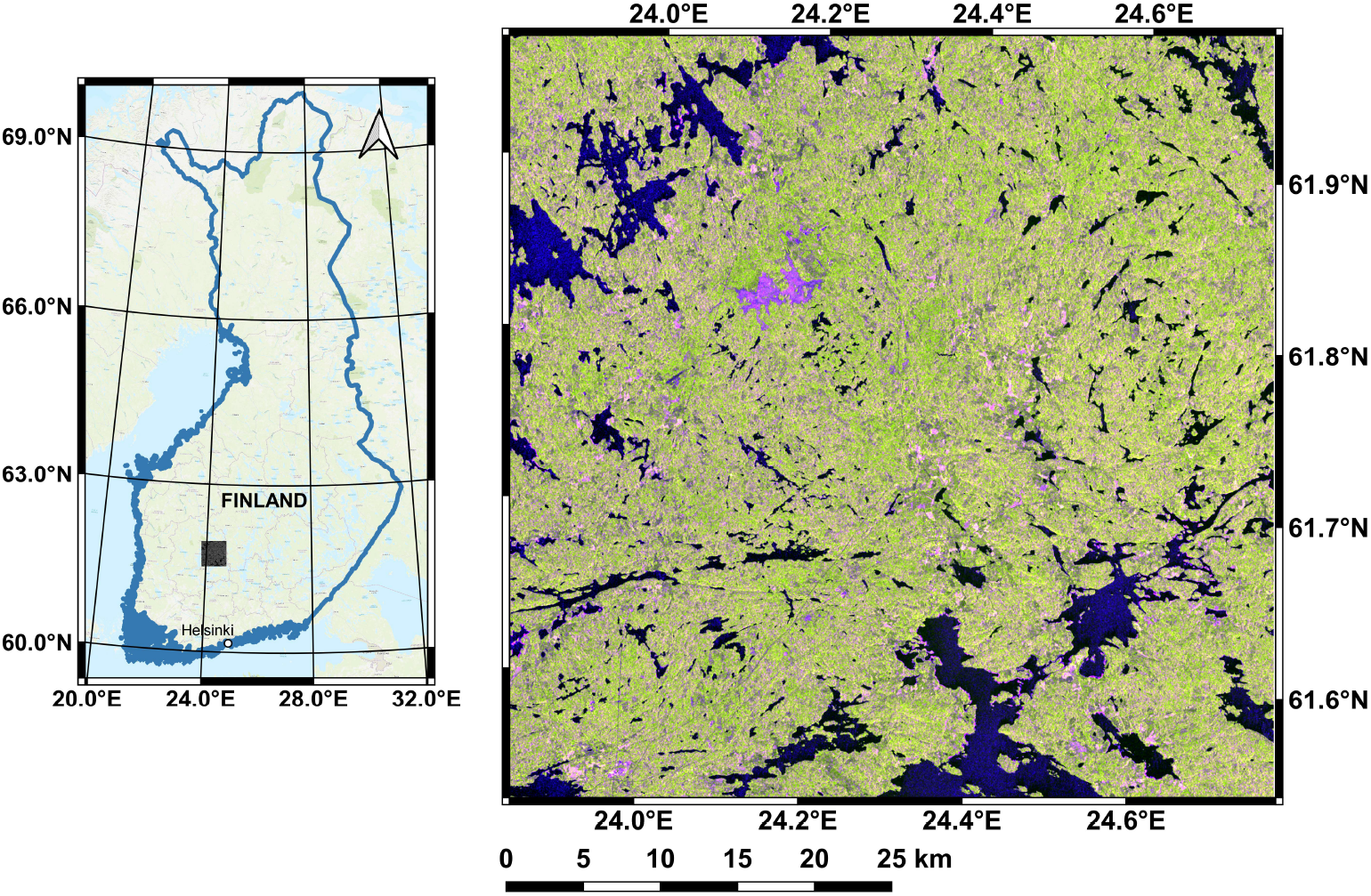
Location of the study site in Southern Finland (left) along with representative Sentinel-1 image as RGB composite (Red: VV, Green: VH, and Blue: VV/VH; Date: 09/10/2014)

### 2.2. Sentinel-1 data

Ninety-six Sentinel-1 images (acquired between 09/10/2014 and 21/05/2018) were used in the study as Level-1 (GRDH, Ground-Range Detected High-resolution) products with VV and VH polarizations. The images were acquired in IW (Interferometric Wide-swath) mode on descending orbits.

The Sentinel-1 images were multilooked with factor 2×2 (range × azimuth) to obtain images with pixel dimensions approximately corresponding to the 20-m grid spacing. The bilinear interpolation method was used for resampling in connection with the ortho-rectification. A digital elevation model (DEM) from the Land Survey of Finland was used and averaged to 20-m × 20-m pixels. Radiometric normalization with respect to the projected area of the scattering element was used to eliminate the topography-induced radiometric variation. In this way, a time series of co-registered images in the terrain corrected “gamma-nought” coefficient was constructed (Small et al., 2010), with a pixel size of 20-m × 20-m. A 3 × 3 median window was also used to filter the speckle noise.

### 2.3. Ground reference data

Stand-level forest resource data that had been collected, prepared and intended for forest management purposes were used as both the training and validation data. The data for the test site were obtained from Finnish Forest Centre (Centre, 2019). These forest resource data were predictions from statistical models using airborne laser scanning (ALS) data and data from field plot measurements. Additionally, field checks on each stand were used to correct noticeable deviations in predictions, e.g., in tree species level predictions, following the practical procedure by the Forest Centre. It is important to keep in mind that despite this procedure, stand-level predictions always include some errors. The data were updated annually based on the reported forestry regimes such as harvests and are thus up-to-date. Because the stand border edges may influence estimation, one-pixel morphological erosion was applied to the stand mask, that is, only pixels separated by at least one pixel from the closest stand boundary were included in the analyses. Further, only stands with areas of at least 1.0 ha after the erosion and with volume of 2.0 m^3^/ha or more were used. The volume threshold was used to remove stands with recent regeneration cuts from the lengthy time series data sets. In total, 17,762 stands were included in our approach of which 8,881 were randomly selected for training and 8,881 remained for validation (Table 1). The stand data were sorted based on the value of the stand ID number. Every second stand was selected for the training data and the rest for the validation data.

**Table 1:** Ground reference data, including the stand area and the growing stock volume, divided into training and validation sets.

| Data set | Number of stands | Stand area, ha |  |  | Volume, m<sup>3</sup>/ha |  |
| --- | --- | --- | --- | --- | --- | --- |
|  |  | Median | Mean | Std | Mean | Std |
| Training | 8,881 | 2.16 | 3.05 | 3.26 | 170.39 | 93.03 |
| Validation | 8,881 | 2.20 | 3.03 | 2.77 | 170.70 | 93.83 |
| Both | 17,762 | 2.20 | 3.04 | 3.03 | 170.55 | 93.43 |

## 3. Methodology

The approach focuses on predicting GSV at stand-level using optional regression techniques, namely multiple linear regression (MLR) (section 3.1.1), support vector machine regression (SVR) (section 3.1.2) and random forests regression (RF) (section 3.1.3).

The general approach is shown schematically in Figure 3. Firstly, a subset of images and one of the optional features were selected. The regression approaches were then used for GSV prediction. This processing was performed on stand-level averaged backscatter (separately for VV- and VH-polarizations, as well as their combination) for selected images to predict stand-level GSV as the target variable. Stand-level averaged backscatter was calculated for each stand using the forest management inventory stand boundaries.

**Figure 2:**
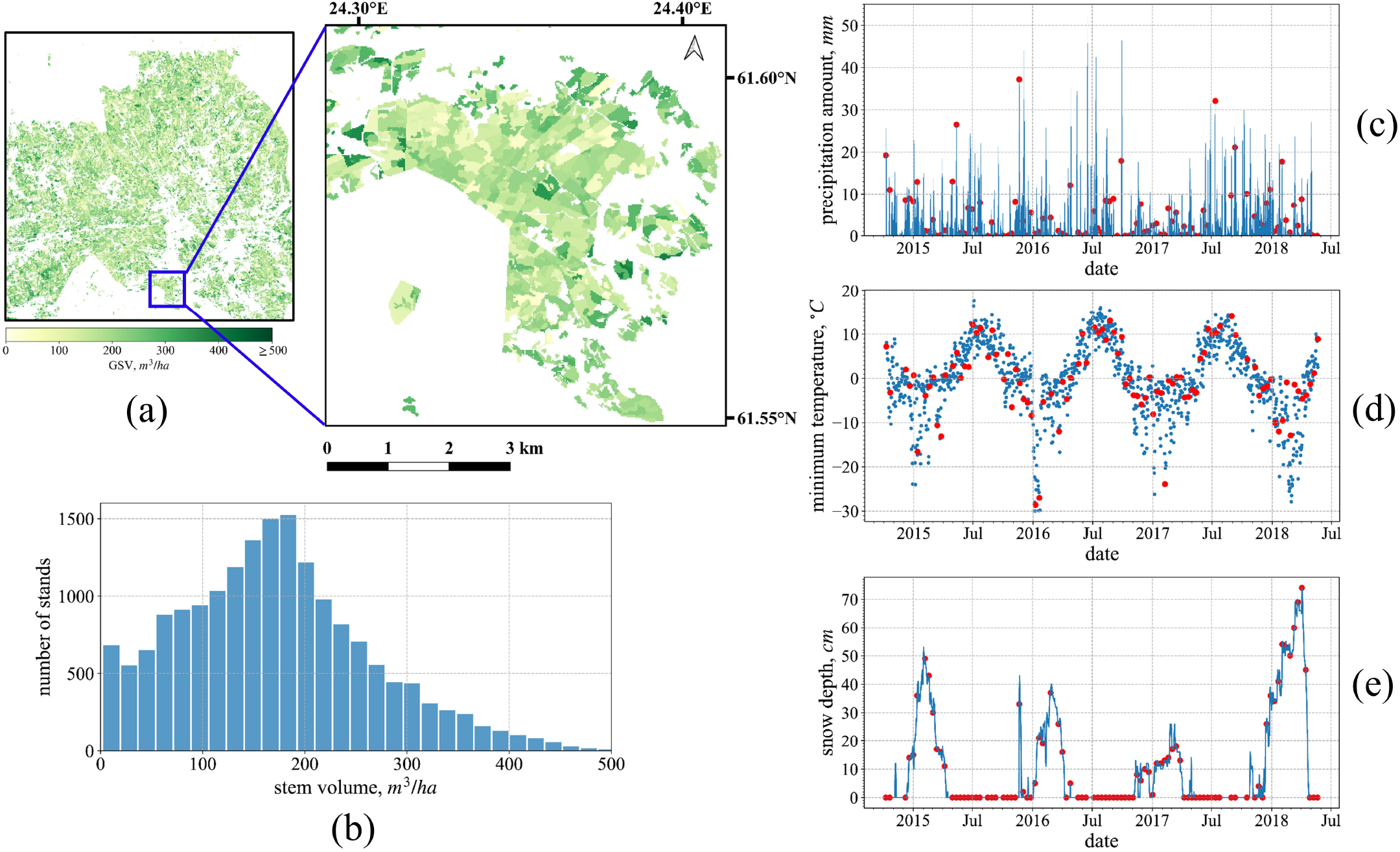
Forest reference data and weather conditions: (a) masks of the forest stands; (b) growing stock volume distribution of all the selected stands for our approaches; (c) three-day accumulated precipitation amount, (d) minimum temperature and (e) snow depth. Sentinel-1 acquisition times are shown with orange marks.

**Figure 3:**
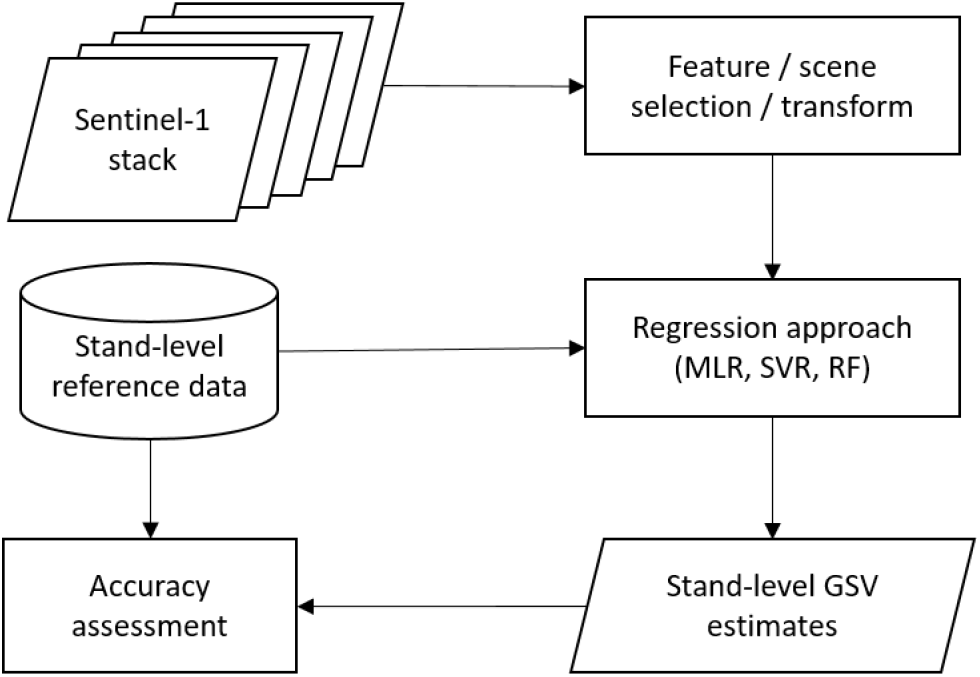
General processing flowchart for growing stock prediction using selected Sentinel-1 images

The stand area after the erosion was used to weight observations, that is stands, in the model estimation, prediction and validation, i.e., RMSEs for the GSV predictions were calculated using Eq. 1

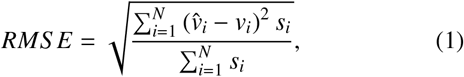

where *v_i_* denotes GSV of stand *i*, 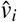 is its prediction, *s_i_* the area of the stand *i* and *N* is the total number of stands.

The GSV prediction approaches used a set of stand-level features as described in section 3.1. The approaches for selecting optimal subsets of features are described in section 3.2.

### 3.1. Methods for predicting growing stock volume

The basic principles of the three prediction methods, MLR, SVR and RF, are described in the following subsections. All methods presume the availability of training data, here stand-level forest resource data.

#### 3.1.1. Multiple Linear Regression

Linear regression was used as one of the parametric methods for predicting GSV. This is a basic regression approach often used for modeling GSV or forest biomass to SAR relationship with various possible transformations of predictor and response variables to linearize the modelled relationship (Dobson et al.,1992; Rauste et al., 1994; Rignot et al., 1994; Kasischke et al.,1995; Englhart et al., 2011; Rauste, 2005; Tsui et al., 2012;Antropov et al., 2013; Hame et al., 2013; Schlund & Davidson,2018; Berninger et al., 2018). We used MLR to be standard for comparisons when using a very long time series of SAR images whose features are used as predictor variables.

Optional transformations of the response variable GSV were tested when selecting the model to predict GSV with an MLR model, including exponential (e.g. with a power of 0.5) and logarithm, in addition to a non-transformed GSV. The final model was of the form

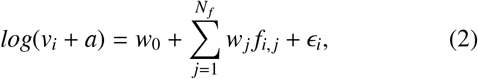

where *v_i_* is GSV for stand *i*, *f_i,j_* the Sentinel-1 feature *j* for stand *i* representing the logarithmic scaled stand-level intensity at VV or VH polarizations or both, *N_f_* the number of the Sentinel-1 features, *W_j_*, j=0,…,N_*f*_ the parameters of the model to be estimated, *a* is a small positive number (0.1) and *∊_i_* random errors assumed to be independently distributed. A bias correction was made to GSV predictions due to the use of a logarithmic transformation in estimating the model Baskerville (1972). The correction was based on Taylor series expansion and assuming that *∊_i_* is normally distributed and is calculated as half of the variance estimate of *∊_i_* (Eq. 2). Note that the correction is additive, is made to the prediction on the log scale before back-transforming, and is half the residual variance on the log scale.

To evaluate the separate contributions to GSV prediction, *f_n_* were selected from different images (timestamps) and polarizations (VV-only, VH-only or VV&VH), as was also the case for other regression approaches described further.

#### 3.1.2. Support Vector Regression

Support vector machines (SVM) classification was developed and first proposed by Vapnik (1995) and Cortes & Vapnik (1995) to solve traditional classification problems. It is based on statistical learning theory. It was extended to the regression problems, support vector regression, SVR, by Vapnik et al. (1997). This approach converts the regression problem to a convex quadratic programming problem. The introduction of the *∊*-insensitive loss ignores the error near true values. SVR uses a non-linear kernel function that transforms a nonlinear problem into a linear one in a higher-dimensional feature space. For the details, see also Gunn et al. (1998); Smola & Schölkopf (2004). The Gaussian Radial Basis function (RBF) kernel was used here. It has shown good performance for forest biomass estimation (Englhart et al., 2011; Neumann et al., 2012). Cross-validation on training data for optimal hyperparameter choices is usually done.

#### 3.1.3. Random Forests Regression

RF is a supervised ensemble learning algorithm that uses a bagging strategy to fit a large collection of classification and regression trees (CART), followed by improving the overall modelling ability Breiman (2001). Since first proposed in 1996, it has been widely used for forest classification, change detection and biomass estimation (Karlson et al., 2015; Belgiu & Drăguţ, 2016; Esteban et al., 2019; Hethcoat et al., 2021)

Compared to SVR or linear regression, RF has multiple advantages: (i) it can avoid overfitting by introducing two-layers of randomisation, i.e., bootstrapping when sampling and random feature selection when splitting nodes (Breiman, 2001); (ii) parallel decision trees not only accelerate the computation but also make nonlinear modelling possible; (iii) high dimensional data can benefit from its automated feature selection, which is especially needed in long time series SAR image processing; and (iv) due to using out-of-bag (OOB) data in internal examination of model, RF can substantially decrease the systematic prediction error without any need for more training data (Gleason & Im, 2012).

In the RF training stage, sub-datasets for each decision tree are randomly selected from the original data using a bootstrapping technique. This ensures the repeatability of training samples for different decision trees. Then, each decision tree is trained independently until each leaf belongs to one specific label. Note that node splitting is based on the features selected as follows. The feature is selected from a random sub-feature space using the criterion characterized as the impurity criterion, which for regression tasks is mean squared error (MSE) or mean absolute error (MAE). In the end, the predictions are averaged to give the final results.

After 5-fold cross-validation on small subsets of training data, the number of trees within the model has been set to 400, and MSE has been chosen as the impurity criterion (variance reduction). To minimise the computational intensity and achieve greater accuracy, the maximum depth of trees varies from 5 to 35, according to the number of input features. The minimum number of sample units required to split an internal node is set to 5 before splitting.

### 3.2. Feature Selection and transformation approaches

Because the Sentinel-1 time series has 96 images covering more than three years, considerable redundancy can be expected in the features derived from them. To circumvent the effects of this redundancy, as well as to reduce the computational burden, feature extraction or reduction can be useful. Here, we describe several approaches for transforming or selecting an optimal subset of SAR features. We discuss both the optimal subset and insights into the choice of the optimal observation date. By carefully choosing a few specific dates, we expect to achieve a prediction accuracy close to what can be achieved using the entire original time series. Principal Component Analysis (PCA) and ik-NN are used for feature transformation and weighting, while feature selection methodologies include a set of techniques based on the ranking strategy: radiometric contrast, mutual information and Lasso. All these approaches are described further.

#### 3.2.1. Principal Component Analysis

Principal Component Analysis (PCA) is a common approach for reducing the dimensionality of feature space (Wold et al.,1987). PCA can be used to convert the original features into a set of linearly uncorrelated components using an eigenvalue decomposition. The importance of each component is evaluated by its eigenvalue. The components corresponding to the largest eigenvalues are expected to retain most information about variation in the data. In this way, a subset of principal components can be used as a set of new features, thereby effectively reducing the original data features while preserving the majority of information. However, because PCA is essentially a feature transformation, and not feature selection approach, it is not suitable for some tasks such as determining the optimal season for acquiring SAR imagery. PCA analysis, therefore, will be primarily used as a standard for comparing feature selection approaches as described further.

#### 3.2.2. Image selection using radiometric contrast

Radiometric contrast is used for choosing a set of images whose feature variables have the greatest radiometric variation between forested and non-forested terrain. This is similar to approaches used for feature selection in earlier studies because of its simplicity and lack of a requirement for reference data (Santoro et al., 2011). The idea is that SAR image features with greater radiometric contrast between forested and non-forested areas would better discriminate between different values of GSV due to a greater dynamic range of forest backscatter measurements.

Forested and non-forested areas are each represented by a specific number of stands that have the least or greatest GSV, respectively. First, we sorted the stands in descending order according to their GSV. Denoting the 10 stands with greatest GSV as the subset **S**_*max*_, and the 10 stands with least GSV as **S**_*min*_ for a specific polarization of a certain image, the radiometric contrast *γ* is evaluated as the ratio of the mean amplitude by using the formula below:

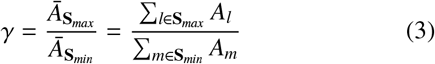

where *A_l_* denotes the stand-averaged feature/backscatter amplitude of stand *l ∈ S_max_*.

Further, all the images were ranked according to the calculated radiometric contrast. The selected images and their relative performance were analysed further in Section 4.2

#### 3.2.3. Feature selection using mutual information

Mutual information is a Shannon entropy-based measure of dependence between random variables. It uses a naive idea that more mutual information means more contributions to regression. Unlike Pearson linear correlation, mutual information can capture nonlinear dependency Kinney & Atwal (2014); Belghazi et al. (2018). The mutual information is computed as

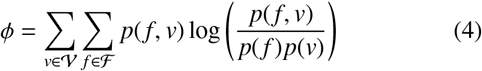

where *f* is the SAR multitemporal feature and *v* is GSV, *p* denotes the probability distribution, *ϕ* is always non-negative and equals zero when and only when 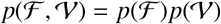.

Accurate estimation of the probability distribution is not trivial, particularly from finite samples. Here we used an approach based on ik-NN to estimate the joint distribution 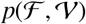 after Kraskov et al. (2004). The number of neighbours, *k*, was set to k=7 in this task (cf. Kinney & Atwal (2014)).

#### 3.2.4. Feature selection using Lasso

Least Absolute Shrinkage and Selection Operator (Lasso) is also a linear regression method. For totals of *N_s_* stands and *N_f_* features, the Lasso loss function is

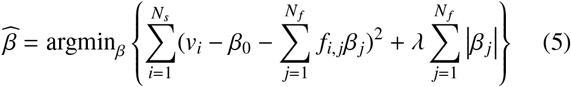

where *f_i,j_* where *f_i,j_* denotes the feature *j* of stand *i*, *V_i_* denotes the corresponding reference GSV, and *β_j_* is the coefficient of feature *j*. The loss is similar to least squares estimation (LSE) for conventional linear regression, the difference is that an L1 regularization 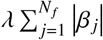 is applied on the coefficients *β_j_* in Lasso. The fitting makes some of the weak feature weights equal to 0, thereby achieving selection of important features. Compared to conventional linear regression, regularization helps to avoid overfitting in our high dimensional condition. *λ* is the weight to control the strength of regularization. Lasso also has some shortcomings. When there is a set of highly relevant variables, Lasso tends to choose one of them and ignores others which may cause instability. In addition, the contribution of each feature is unclear because of the sparsity of features leading to difficulties in understanding the data.

#### 3.2.5. Feature selection using a genetic algorithm

The well-known k-NN estimation method is well-suited for cases in which multiple target parameters or variables are estimated simultaneously. In this paper, it was tested as an optional method for feature weighting and selection. The weights for the features selected were calculated as per Tomppo & Halme (2004); Tomppo et al. (2008a,b, 2019).

Further we briefly recall the main properties of the ik-NN approach with the genetic algorithm in the context of feature selection. Let us consider a stand *p* and denote its *k* nearest feasible field stands by *i*_1_(*p*),…,*i_k_*(*p*) when the distance is measured in the feature space. The weight *w_i,p_* of field stand *i* to stand *p* is defined as

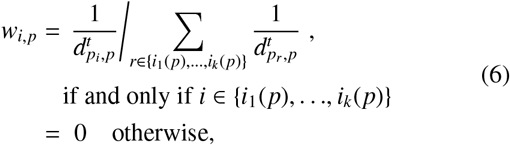

where *p_r_* is a stand with an index *r* and *r* belongs to the set {*i*_1_(*p*),…,*i_k_*(*p*)}. The value of *k* was fixed to be 15 after tests using the RMSEs and mean deviations in the validation data set as the criteria. The distance weighting power *t* is a real number, usually *t* ∈ [0,2]. The value *t*=1 was used here. A small quantity, greater than zero, is added to *d* when *d* = 0 and *i* ∈ {*i*_1_(*p*),…*i*_k_(*p*)}.

The distance metric *d* employed was

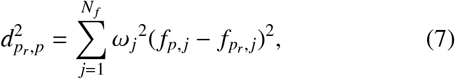

where *f_p,j_* is the *j*-th SAR feature variable for stand *p*, *N_f_* the number of SAR feature variables, and *ω_j_* the weight for the *j*-th SAR feature variable.

The values of the elements *ω_j_* of the weight vector *ω* were selected with a genetic algorithm (Tomppo & Halme, 2004; Tomppo et al., 2008a).

The fitness function to be minimized for calculating the values of the *ω* vector for GSV estimation was

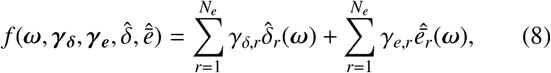

where

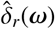 is the standard error of the prediction of the forest variable *i* at stand-level,
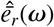 is the mean deviation of the forest variable *i* at stand-level,
*N_e_* the number of the forest variables used in the algorithm (here one) and
***γ_δ_*** and ***γ_e_*** are fixed constant vectors.

After some trial and error tests and heuristic trade-offs between standard error and mean deviation, the values of ***γ_δ_*** = 10 and ***γ_e_*** = 1 were selected, thus giving RMSE greater weight. Otherwise, the parameters of the genetic algorithm itself were the same as in Tomppo & Halme (2004). Other parameters are, e.g., 1) the number of generations, 2) the number of weight vectors in one population and in the medipopulation, 3) the probabilities of uniform crossover, accepting an inferior solution created by mutation, mutation, and radical mutation. For the details, see Tomppo & Halme (2004). As an iterative procedure, genetic algorithms are computationally intensive and may produce a local rather than global optimum. To circumvent this possibility, multiple runs are usually needed to find a near global optimum.

The weights were sought for each combination of images and polarizations separately.

### 3.3. Multitemporal analysis

Multitemporal analysis of SAR imagery for this study included evaluation of accuracy gain achieved using multiple SAR images. These images were used either as sequences of consecutively collected images, or specific images as chosen using feature selection approaches (section 3.2). For the latter, optimal SAR images can be spread over the 3-year observation period. However, using all data seems impractical, because strong saturation can be observed and any additional value of using more than 10 images can be questionable. Also, use of such a long time series can be computationally very demanding. Thus, selection of optimal timing for collecting imagery should be attempted using RMSE as the selection criterion. To determine the optimal timing, that is image acquisition season, and to analyse the “seasonality” for collecting SAR imagery, a “sliding window” analysis was performed.

#### 3.3.1. Rationale for “sliding window” analysis

The size of the sliding window corresponds to the number of images that are aggregated and from which the feature variables are derived. Several aggregation approaches are possible in the context of GSV estimation. The first, multitemporal filtering, facilitates reduction of speckle by calculating the combined estimate from several images via averaging (Quegan et al., 2000). Filtered image is further used for calculating image features for predicting GSV. Another approach performs image-by-image GSV estimation followed by merging individual image-based estimates using a multiple regression approach (Kurvonen et al.,1999). Individual GSV estimates can be obtained using methods such as inversion of physics-based models (Antropov et al.,2013; Santoro et al., 2021). However, for this study, a set of feature variables derived from SAR images can be used directly as input to a multiple regression approach, thereby constituting a third possible approach. It seems also most practical.

Thus, the prediction takes the form:

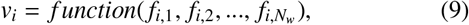

where *N_w_* is the size of the sliding window, and *i* is the stand number.

The choice of the size of the sliding window, i.e., the number of images from which the feature variables were selected for the regression, thereby requires separate experimentation. On one hand, including more images is expected to improve GSV prediction performance. On the other hand, the value of additional images can be limited as the number of images increases beyond a minimal number, not to mention the increased computational intensity associated with larger sliding windows. Thus, several window sizes from 1 to 32 were tested for this study.

## 4. Results

We started our analyses by assessing the improvement in GSV prediction as the number of Sentinel-1 images increases when using the three regression approaches. The effect on the image acquisition time on the prediction and relative performance of the regression approaches are reported in section 4.1.

Images are aggregated in the order of their acquisition. Further, the potential of feature selection and transformation approaches and their effect on the accuracy of GSV prediction with the regression approaches are analysed in section 4.2. The effects of combining a small number of images within the “sliding window”for GSV prediction are reported in detail in section 4.3.

### 4.1. Accuracy of GSV prediction

#### 4.1.1. Single SAR images and seasonal variation of the predictions

Figure 4 shows the accuracy of the GSV predictions when using feature variables derived from single SAR images acquired from Nov 2014 to July 2018. The accuracy metrics are shown in the order of image acquisition. RMSE varies within the range 85-94 m^3^/ha (rRMSE 50-55%) for various polarizations. Overall, the dynamics were quite variable as expected for C-band imagery which is strongly affected by environmental conditions. For MLR, the greatest accuracies were obtained using feature variables selected for images for the month of June for 2015, 2016 and 2017. For SVR and RF, a similar observation can be made with an optimum in early 2016. For all image and polarization combinations, the combined VV and VH polarizations gave smaller RMSEs than any single polarization alone, and VV-pol somewhat smaller than VH-pol. Scatterplots of RMSEs using the different GSV prediction methods for several selected sets of Sentinel-1 image features are shown in Fig. 5.

**Figure 4:**
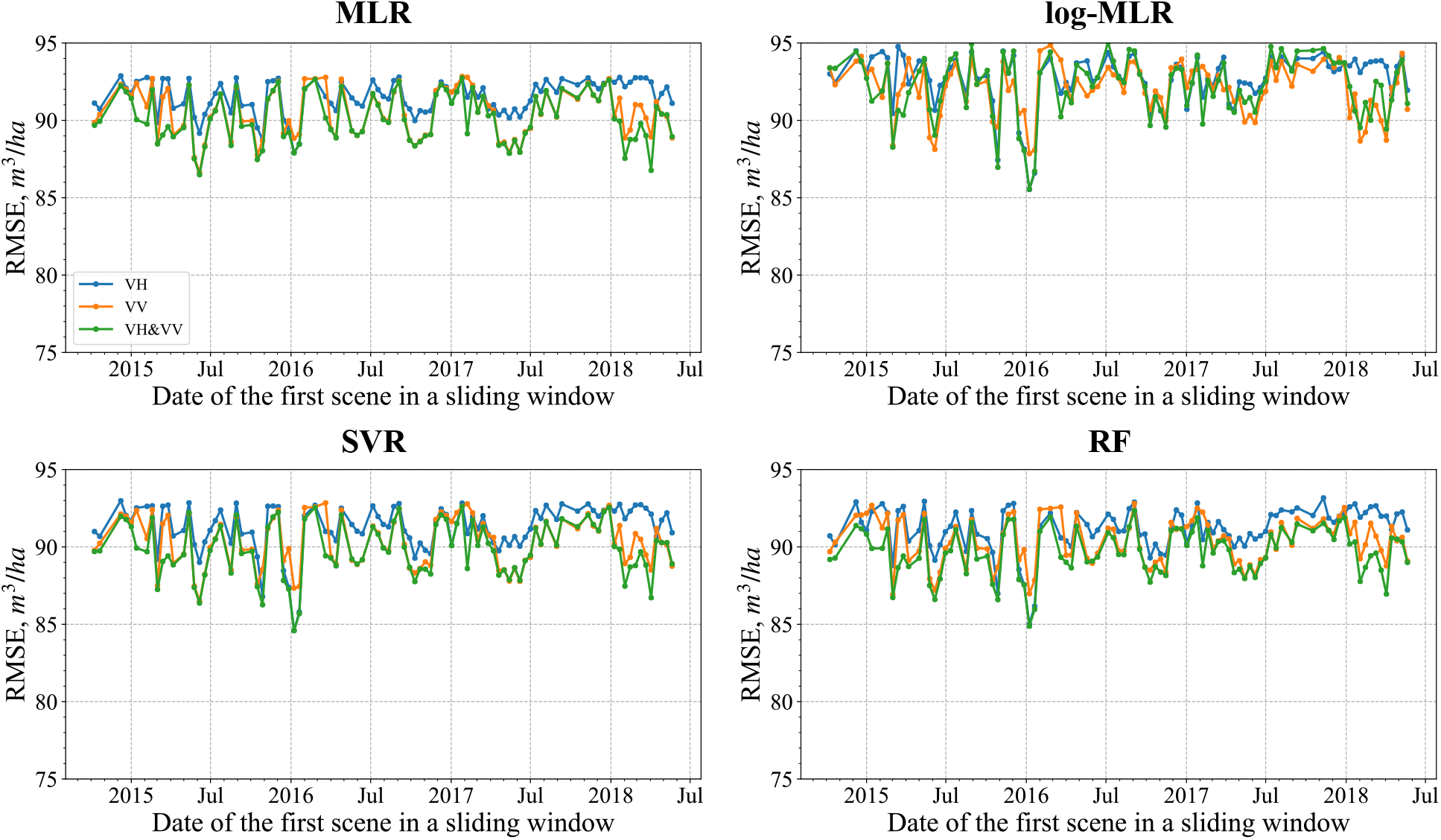
RMSEs of the predicted GSV using individual images (both polarizations) with the three methods MLR, log-MLR, SVR and RF

**Figure 5:**
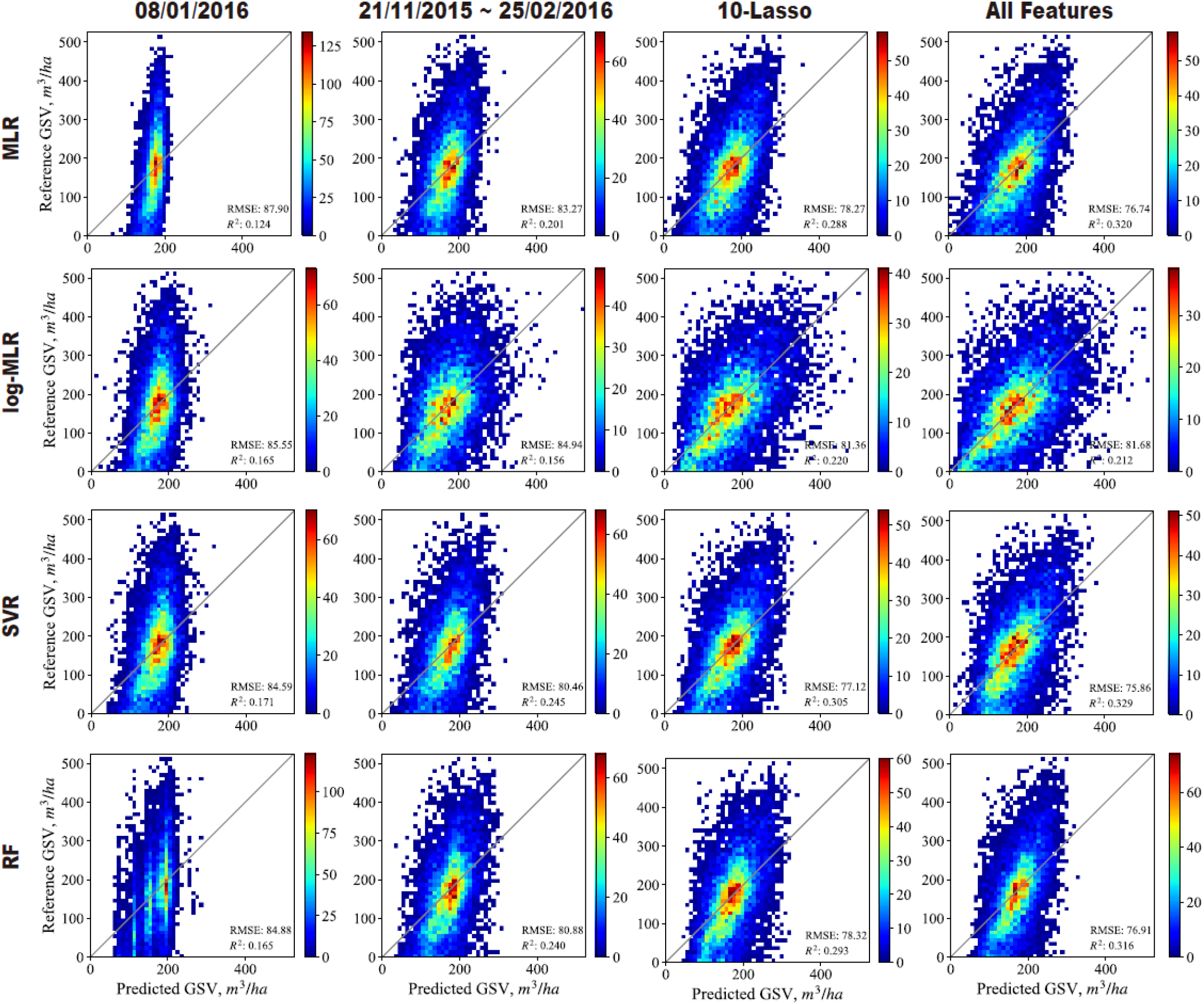
Scatterplots of the final estimation. Each row corresponds to MLR, log-MLR, SVR and RF, separately. First column is regression results on only one single image (08/01/2016), second column is regression results on eight consecutive scenes (21/11/2015 25/02/2016), third column is regression results on all 192 features, the last column is regression results on 10 best Lasso features.

#### 4.1.2. Accumulated SAR time series

We further tested the accuracy of the GSV predictions by using feature variables for multiple images (SAR time series) in the modelling with the three different methods, MLR, SVR and RF and with three polarization options VH, VV and both VV and VH. The variation in RMSEs for the GSV predictions are illustrated in Fig. 6. Here, the images were accumulated in the order of their acquisition. GSV prediction for any given date is performed using all images acquired until the date on x-axis. Altogether nine RMSEs as a function of the number of accumulated images used in the GSV prediction are presented in Fig. 6.

**Figure 6:**
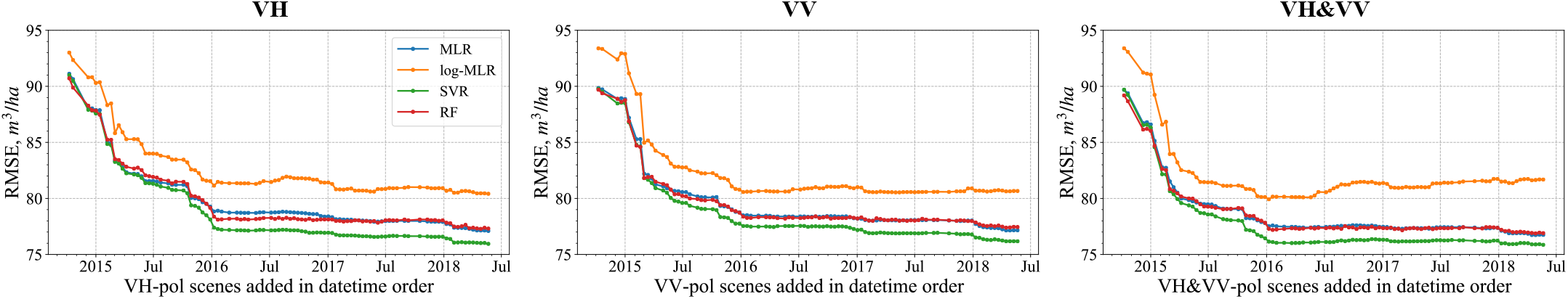
Accuracy of predicted GSV with image features in the order of image acquisition.

A nearly monotonic decrease in RMSEs for the GSV predictions for all the polarization combinations and all three prediction methods can be seen. The RMSEs rapidly decreased initially but levelled off shortly after July 2015 by which time more than 15 images had accumulated in the SAR time series. RMSE decreases somewhat faster for VV-pol than for VH-pol, with RMSEs approximately 80 m^3^/ha for VV and approximately 82 m^3^/ha for VH for July 2015. Combined VH and VV gave smaller RMSE for GSV prediction than VV or VH alone, less than 76 m^3^/ha (rRMSE 45%) for the same date.

For the majority of our experiments, that is, combinations of image sets, polarization and prediction technique, SVR produced somewhat smaller RMSEs than MLR or RF, about 1.0 m^3^/ha smaller, whereas there was practically no difference between the RMSEs for MLR and RF, except for a short period from 2016 to 2017 for VH-pol.

### 4.2. Feature Selection

Here, we evaluated the potential of Sentinel-1 for GSV prediction using selected sets of images in contrast to using images in the order of their acquisition date as in section 4.1.2. The idea was to achieve the greatest GSV prediction accuracy using a minimum number of images. In the further analyses, we evaluated several feature selection approaches. When particular images were selected, their features were used in stand-level GSV prediction.

The results of GSV predictions using different feature selections and transformation approaches to form multitemporal SAR stacks are presented in Figure 7. Both polarizations were used together, thus the size of the original feature pool equals 192. The PCA results are also shown in Figure 7 as a benchmark. In the figures, the curves for only the first 50 components are shown, due to the lack of accuracy improvement after this number.

**Figure 7:**
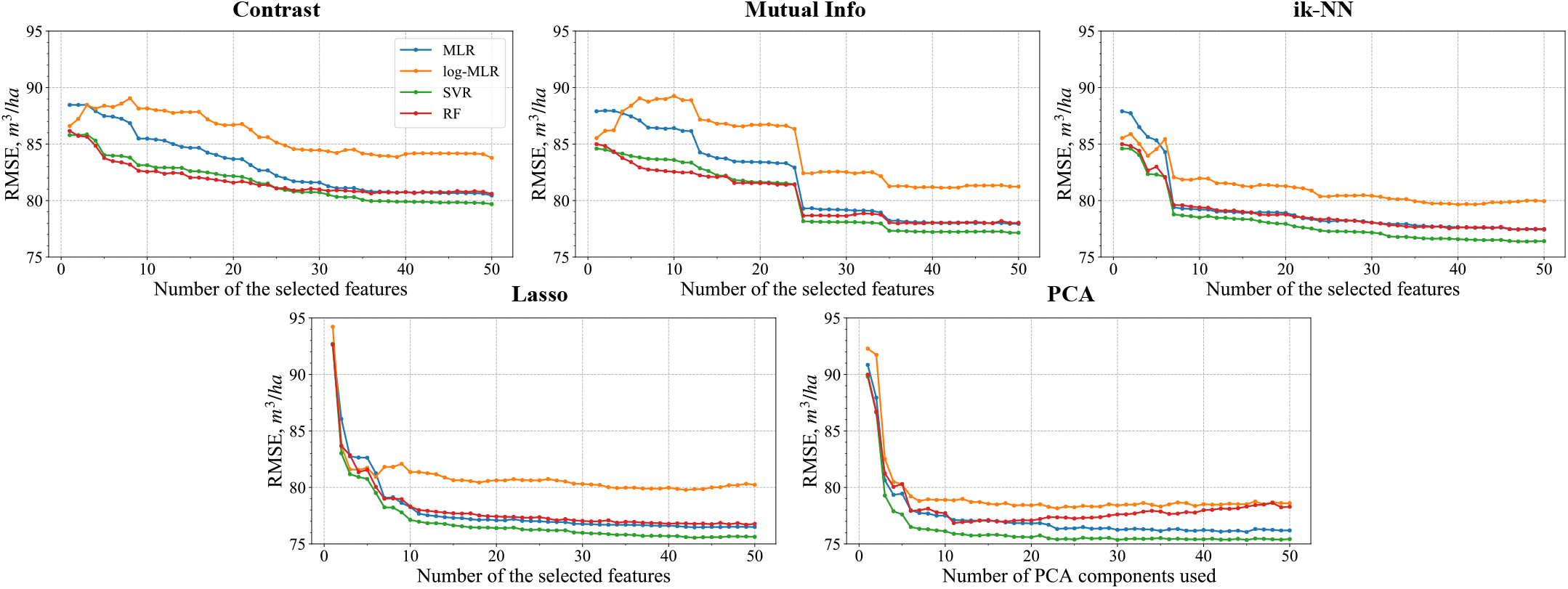
RMSE against feature streams. All 192 features (VH+VV) are ranked, first 40 are shown in the figure. Features are ranked using five methods: Radiometric Contrast, Mutual Information, Lasso, ik-NN and PCA respectively. All four regressions are used in the evaluation.

The SVR method produced slightly smaller RMSEs than MLR or RF, similar to the results in section 4.1.2 (Fig. 6). RF produced somewhat smaller RMSEs than MLR. Especially, MLR gave larger RMSEs when using the initial 20-25 features and the features selected using radiometric contrast and mutual information. All three prediction methods produced smaller RMSEs when using ik-NN, Lasso and PCA feature selections than with the two other feature selection methods, radiometric contrast and mutual information. The convergence to maximum accuracy levels was much slower with the latter methods. This indicates that a blind feature selection approach (with no reference data) such as radiometric contrast can be considerably sub-optimal. However, it is the only possible approach in the absence of reference data. It is also notable that when using several initial images with high-contrast features, RMSEs were smaller than when the images were used in order of acquisition date. The convergence to the maximum accuracy was much slower in the latter case.

Interestingly, RF-based RMSEs for GSV with PCA-selected features started to increase when more than 20 selected principle components were included in the model. A likely reason is that the PCA tail-components consist largely of noise because such phenomena were not detected with RF when using the backscattering coefficients of the original or selected images in GSV prediction. Other approaches, MLR and SVR, appeared more robust in this respect, with consistent decreases even when incorporating noise-alike PCA features. The achieved “plateau” of RMSE was approximately 75.1 m^3^/ha (rRMSE 44%).

While the feature selection approaches are based on various ranking principles, one possible way to compare their performance is to evaluate how “quickly” the gain in GSV estimation accuracy is achieved when the selected features are combined. An approach that delivers a quick and stable decrease on GSV prediction RMSE can be recommended as an optimal feature selection approach in the context of this study.

Among all feature selection approaches, Lasso produced the smallest RMSE values and the greatest rate of RMSE decrease when the additional features were added, indicating that it might be the most effective approach for selecting several variables. RMSEs for all the regressions achieved a level of less than 78 m^3^/ha when only 20 selected features were used. The fact that only 20 features were enough to obtain large prediction accuracies is similar to observations made with PCA-derived features. Similarly, the second-best performing approach, ik-NN achieved 79 m^3^/ha. Radiometric contrast appeared least capable as a method for selecting features because of the relatively slow rate of decrease of RMSE as the number of features increased. RMSE-dependence for mutual information feature selection approach exhibited a “step-wise” decrease, indicating this feature selection approach was also suboptimal because the features that most increased accuracy were not included early enough.

Similar observation can be made from Figure 5. Here, SVR produced scatter graphs indicating greater accuracy in terms of both RMSE and R^2^. The feature selection approaches providing greatest GSV prediction accuracies are illustrated as “10-PCA” and “10-Lasso”, respectively.

When SVR was used (see Figure 8), the effect of polarization was small, particularly for PCA and Lasso. Radiometric contrast and mutual information were the least accurate among feature selection approaches, indicating that the simplest approaches are not necessarily most useful. Among these two approaches, mutual information was still more accurate when VV-pol and combined VV and VH were used, while there was no difference for VH-pol.

**Figure 8:**
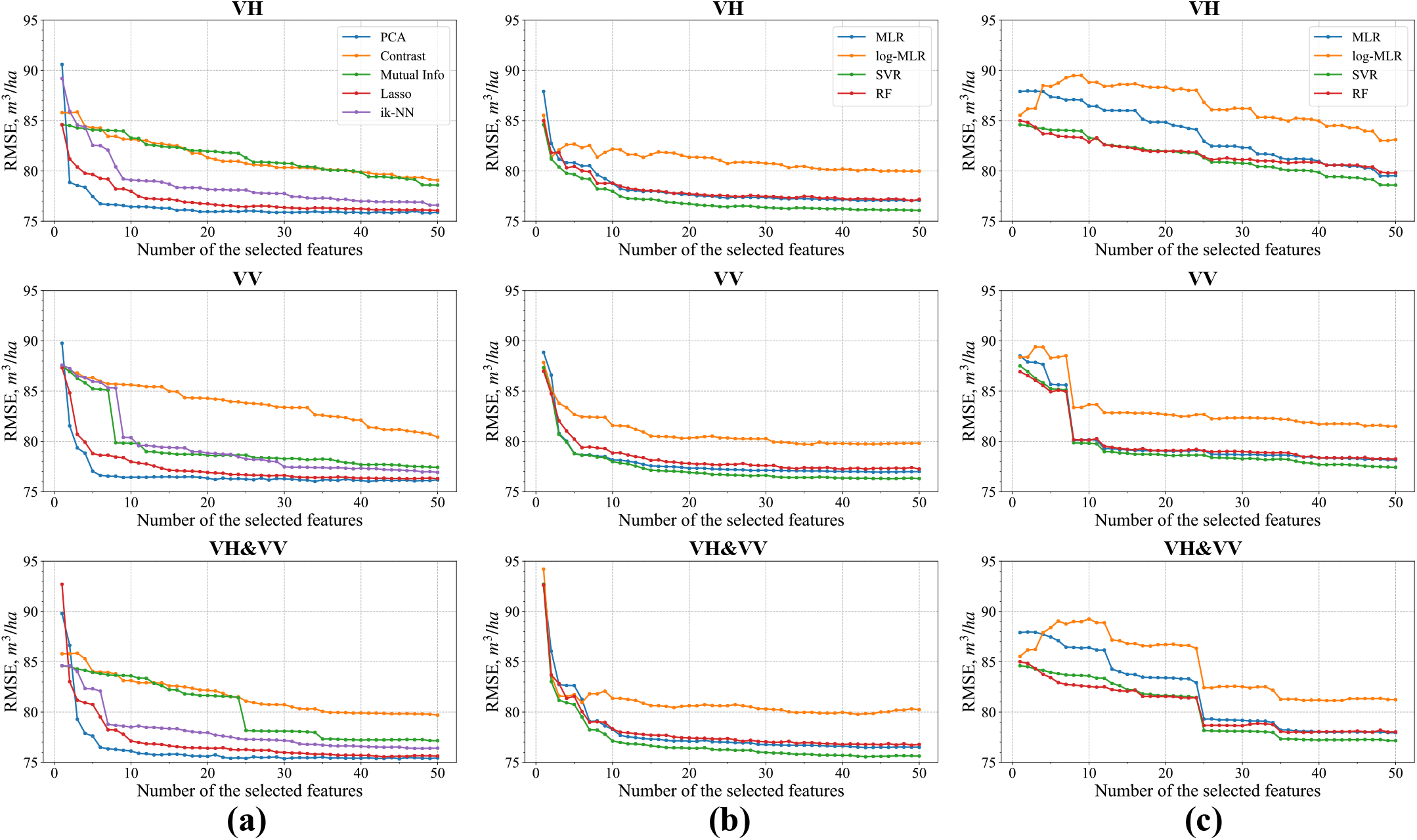
RMSE dependence on the number of input features at various polarizations: (a) all feature selection approaches with SVR prediction; (b) Lasso feature selection approach with all regression approaches; (c) Mutual Information feature selection approach with all regression approaches

MLR performed a bit less accurately than the two other approaches for GSV prediction when mutual information was used for the feature selection (Figure 8). All prediction methods behaved practically in the same way with Lasso-based features. Mutual information-selected features with combined (VV and VH) image stack performed even less accurately than VV-only images with MLR. That was not the case for SVR and RF. In contrast, the Lasso approach for the combined feature set (VH and VV) achieved greater accuracies compared to either VH-only or VV-only feature selection. The final scatter graphs (Figure 5) for “10-Lasso” and “10-PCA” both showed very similar dependence and accuracies when compared to “All Features”; this further confirmed the efficiency of the suggested Lasso approach for SAR feature selection.

ik-NN was the second-best approach with respect to GSV prediction accuracy in practically all experiments. For ik-NN selected features, the GSV prediction accuracy dependence for VH-pol decreased faster than for VV-pol for the first 10 selected features (see Figure 8).

Overall, Lasso performed most accurately among all the feature selections, consistently providing greatest GSV accuracy with all regression methods. VH-pol appeared consistently close to VV-pol for selected features (except for mutual information). SVR was confirmed to be the most accurate method in feature based GSV prediction.

Table 2 shows feature selection results for the three polarization combinations. The key features (last column within tables) were chosen using a simple voting strategy according to image ranking by all feature selection approaches. When a specific image ranked first, it got 10 points, 2nd best scores 9 and so on. After adding all the points for a given image, it got its final rank in the “key feature” column. Noteworthy, in the ranking scenario where both polarizations were mixed, the ranking order delivered by Lasso and ik-NN may be different compared to cases when only VH- or VV-pol bands were considered.

**Table 2:**
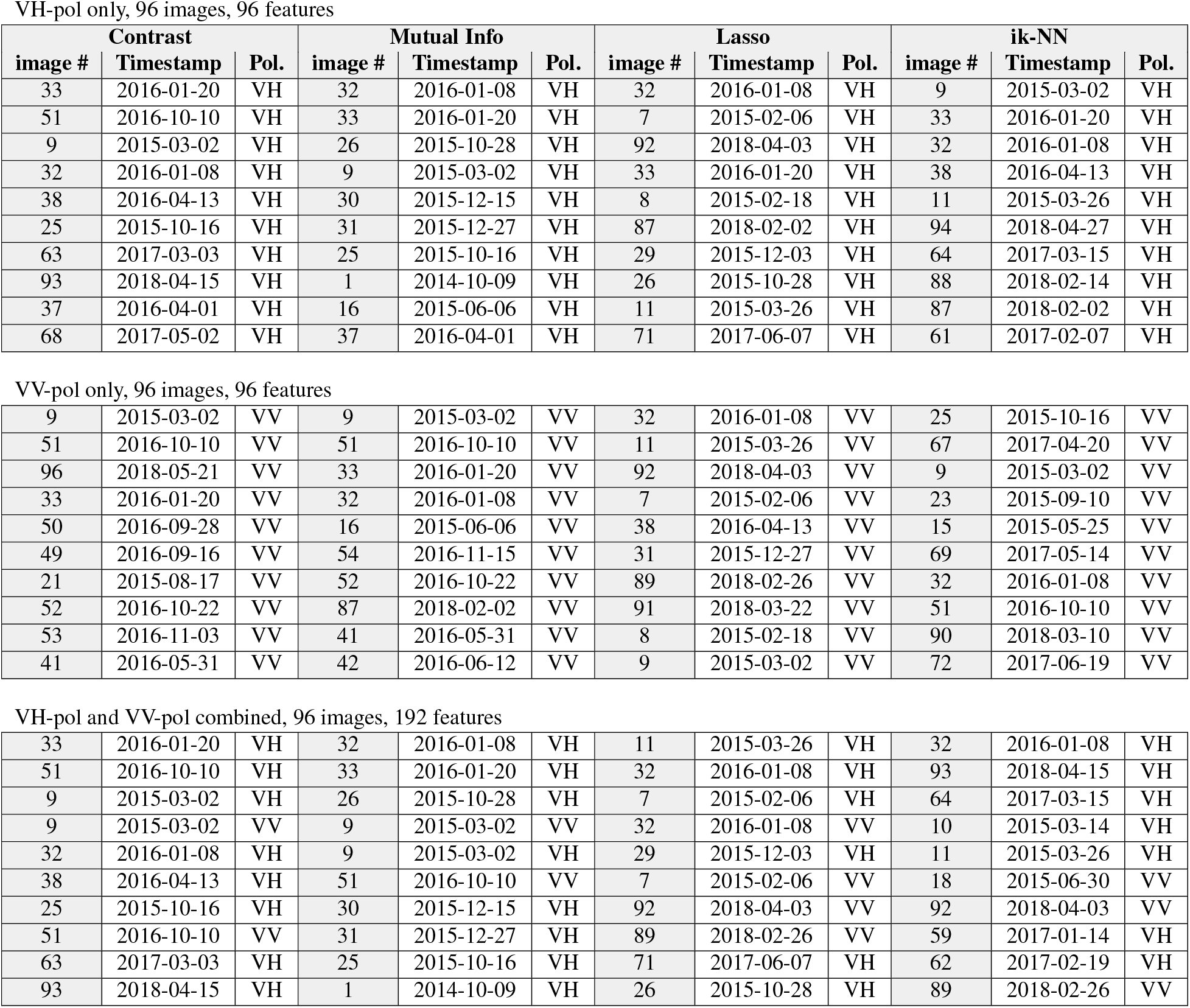
Feature selection results using several feature selection approaches and majority voting

VH-pol only, 96 images, 96 features
| Contrast |  |  | Mutual Info |  |  | Lasso |  |  | ik-NN |  |  |
| --- | --- | --- | --- | --- | --- | --- | --- | --- | --- | --- | --- |
| image # | Timestamp | Pol. | image # | Timestamp | Pol. | image # | Timestamp | Pol. | image # | Timestamp | Pol. |
| 33 | 2016-01-20 | VH | 32 | 2016-01-08 | VH | 32 | 2016-01-08 | VH | 9 | 2015-03-02 | VH |
| 51 | 2016-10-10 | VH | 33 | 2016-01-20 | VH | 7 | 2015-02-06 | VH | 33 | 2016-01-20 | VH |
| 9 | 2015-03-02 | VH | 26 | 2015-10-28 | VH | 92 | 2018-04-03 | VH | 32 | 2016-01-08 | VH |
| 32 | 2016-01-08 | VH | 9 | 2015-03-02 | VH | 33 | 2016-01-20 | VH | 38 | 2016-04-13 | VH |
| 38 | 2016-04-13 | VH | 30 | 2015-12-15 | VH | 8 | 2015-02-18 | VH | 11 | 2015-03-26 | VH |
| 25 | 2015-10-16 | VH | 31 | 2015-12-27 | VH | 87 | 2018-02-02 | VH | 94 | 2018-04-27 | VH |
| 63 | 2017-03-03 | VH | 25 | 2015-10-16 | VH | 29 | 2015-12-03 | VH | 64 | 2017-03-15 | VH |
| 93 | 2018-04-15 | VH | 1 | 2014-10-09 | VH | 26 | 2015-10-28 | VH | 88 | 2018-02-14 | VH |
| 37 | 2016-04-01 | VH | 16 | 2015-06-06 | VH | 11 | 2015-03-26 | VH | 87 | 2018-02-02 | VH |
| 68 | 2017-05-02 | VH | 37 | 2016-04-01 | VH | 71 | 2017-06-07 | VH | 61 | 2017-02-07 | VH |
| VV-pol only, 96 images, 96 features |  |  |  |  |  |  |  |  |  |  |  |
| 9 | 2015-03-02 | VV | 9 | 2015-03-02 | VV | 32 | 2016-01-08 | VV | 25 | 2015-10-16 | VV |
| 51 | 2016-10-10 | VV | 51 | 2016-10-10 | VV | 11 | 2015-03-26 | VV | 67 | 2017-04-20 | VV |
| 96 | 2018-05-21 | VV | 33 | 2016-01-20 | VV | 92 | 2018-04-03 | VV | 9 | 2015-03-02 | VV |
| 33 | 2016-01-20 | VV | 32 | 2016-01-08 | VV | 7 | 2015-02-06 | VV | 23 | 2015-09-10 | VV |
| 50 | 2016-09-28 | VV | 16 | 2015-06-06 | VV | 38 | 2016-04-13 | VV | 15 | 2015-05-25 | VV |
| 49 | 2016-09-16 | VV | 54 | 2016-11-15 | VV | 31 | 2015-12-27 | VV | 69 | 2017-05-14 | VV |
| 21 | 2015-08-17 | VV | 52 | 2016-10-22 | VV | 89 | 2018-02-26 | VV | 32 | 2016-01-08 | VV |
| 52 | 2016-10-22 | VV | 87 | 2018-02-02 | VV | 91 | 2018-03-22 | VV | 51 | 2016-10-10 | VV |
| 53 | 2016-11-03 | VV | 41 | 2016-05-31 | VV | 8 | 2015-02-18 | VV | 90 | 2018-03-10 | VV |
| 41 | 2016-05-31 | VV | 42 | 2016-06-12 | VV | 9 | 2015-03-02 | VV | 72 | 2017-06-19 | VV |
| VH-pol and VV-pol combined, 96 images, 192 features |  |  |  |  |  |  |  |  |  |  |  |
| 33 | 2016-01-20 | VH | 32 | 2016-01-08 | VH | 11 | 2015-03-26 | VH | 32 | 2016-01-08 | VH |
| 51 | 2016-10-10 | VH | 33 | 2016-01-20 | VH | 32 | 2016-01-08 | VH | 93 | 2018-04-15 | VH |
| 9 | 2015-03-02 | VH | 26 | 2015-10-28 | VH | 7 | 2015-02-06 | VH | 64 | 2017-03-15 | VH |
| 9 | 2015-03-02 | VV | 9 | 2015-03-02 | VV | 32 | 2016-01-08 | VV | 10 | 2015-03-14 | VH |
| 32 | 2016-01-08 | VH | 9 | 2015-03-02 | VH | 29 | 2015-12-03 | VH | 11 | 2015-03-26 | VH |
| 38 | 2016-04-13 | VH | 51 | 2016-10-10 | VV | 7 | 2015-02-06 | VV | 18 | 2015-06-30 | VV |
| 25 | 2015-10-16 | VH | 30 | 2015-12-15 | VH | 92 | 2018-04-03 | VV | 92 | 2018-04-03 | VV |
| 51 | 2016-10-10 | VV | 31 | 2015-12-27 | VH | 89 | 2018-02-26 | VV | 59 | 2017-01-14 | VH |
| 63 | 2017-03-03 | VH | 25 | 2015-10-16 | VH | 71 | 2017-06-07 | VH | 62 | 2017-02-19 | VH |
| 93 | 2018-04-15 | VH | 1 | 2014-10-09 | VH | 26 | 2015-10-28 | VH | 89 | 2018-02-26 | VV |

For both VV and VH, key features belonged mostly to images acquired under frozen conditions. However, several “dry summer” images acquired in May and June were also included by most accurate Lasso and ik-NN approaches. We performed additional experiments by Monte Carlo testing of random combinations of 10 images representing frozen ground conditions (VV and VH features combined) in GSV prediction using SVR approach. After 50 trials, RMSE was 79.3 ±1.1 m^3^/ha and never exceeded the accuracy of the “10-Lasso” approach of 77.1 m^3^/ha when all images were used (Table 2). Corresponding accuracy for 10 ik-NN selected features using all possible images reached 78.5 m^3^/ha exceeding absolute majority of trials with 10 exclusively frozen condition images. Similar dynamics were observed also for other regression approaches, indicating that the idea of using dry summer images in addition to frozen condition images should be examined in further studies. Cross-pol performed slightly more accurately than co-pol. Several images, such as those acquired 2015/03/02 and 2016/01/08, had both polarizations among selected features, indicating that nearly optimal GSV prediction can be achieved using only a few carefully selected images (instead of the whole SAR image stack). This observation is in line with PCA feature selection discussed above.

### 4.3. Sliding Window Analysis

The sliding window analyses started by selecting a suitable window size for the further tests (section 3.3.1). For this purpose, we used SVR as the prediction method and tested several window sizes: from 1, corresponding to one image only, to as many as 15 images.

The results of GSV prediction using sliding windows of different sizes are shown in Figure 9. The date of the first image within a sliding window is shown on the x-axis. RMSE varied from 85.3 m^3^/ha to 93.5 *m*^3^/*ha* with an average RMSE of ~90 m^3^/ha when the sliding window size *N_w_* equals 1, that is, only one image at a time was used to predict GSV. The greatest accuracy was achieved using the image acquired on Jan. 08, 2016. No pronounced RMSE seasonal dynamics can be observed. Further, as *N_w_* gets larger, the RMSE decreased gradually. The largest RMSE with *N_w_*=4 was ~92 m^3^/ha. The largest RMSE became smaller than 88 m^3^/ha with *N_w_*=15.

**Figure 9:**
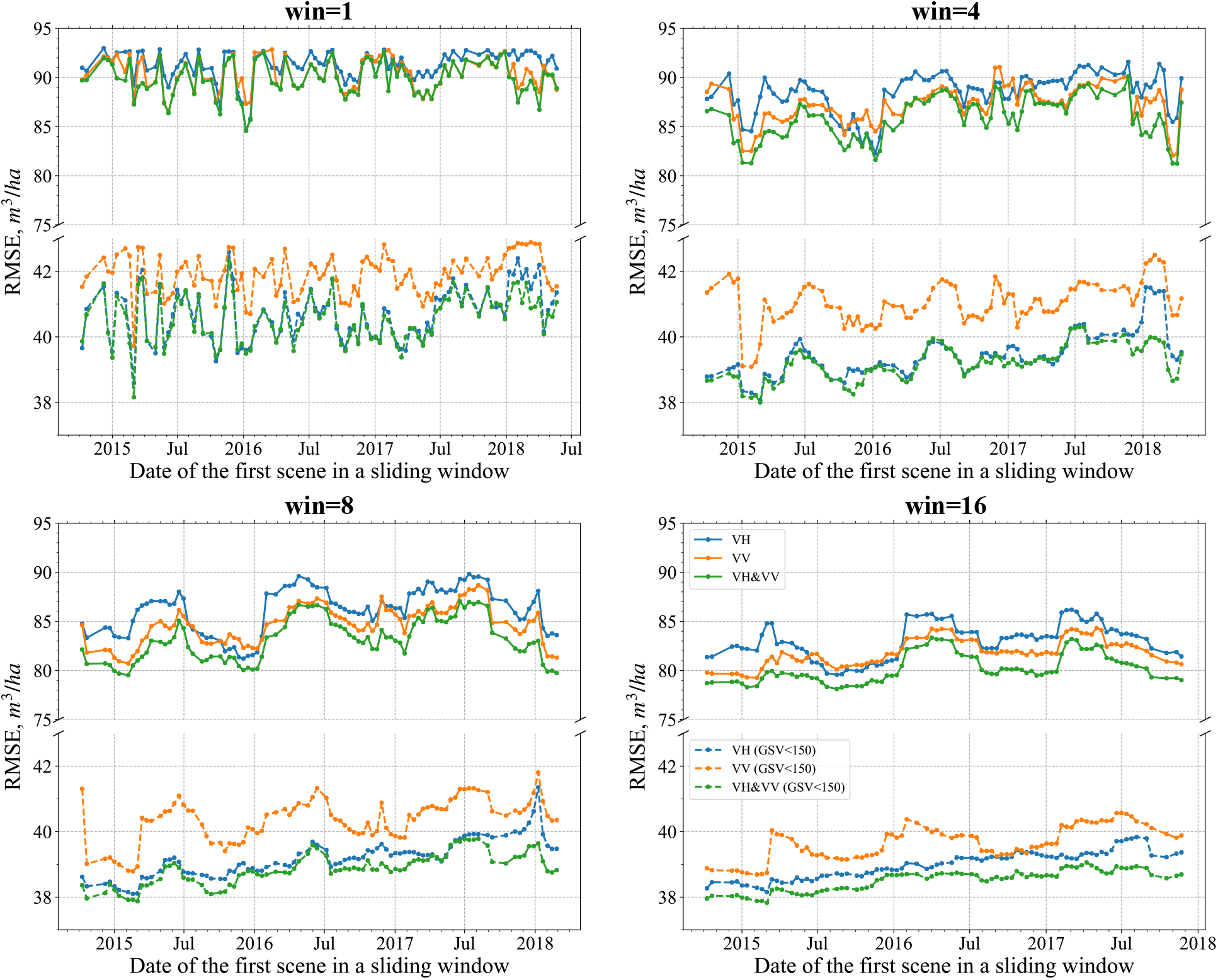
RMSE of GSV prediction as a function of image acquisition date for various sliding windows. SVR approach is used. The window sizes are 1, 4, 8 and 16. Solid curves show RMSE show experiments with all stands, and dashed curves show RMSE results by using stands whose biomass are less than 150*m*^3^/*ha*

The seasonal patterns, indicated by the RMSE curves, became apparent and more pronounced when expanding the sliding window size. For example, in Figure 9(b), the seasonal pattern is not very clear, whereas in Figure 9(d) we can readily observe annual variations: the local minima of RMSE-curves are clearly visible in every winter, while the local maxima can be found every summer. When the window size is very large, approaching six months, the second peaks are almost eliminated and the three main annual peaks become clearly visible (Figure 9(f)).

Note that the x-axis tick marks denote acquisition dates for the first Sentinel-1 image within each sliding window, whereas all images within the window contribute to GSV prediction. The temporal resolution decreases when the window size increases, with the RMSE temporal variation becoming smooth. The width of the extreme is particularly affected. Consider the situation around January 2016 as an example. When the window size is small, 1-4, the temporal fluctuations in RMSE, that is, decreases and increases, are rapid, and the valleys are sharp, as highlighted by the red circles in Figure 9(a). When the window size increases, the fluctuation becomes smoother. The temporal range of extreme exceeds half a year in Figure 9(f). Similarly, the widths of major peaks are smoothed. The reason is that the image on Jan/08/2016 contributed strongly to the GSV prediction with all sliding windows (that include Jan/08/2016) exhibiting large prediction accuracies. This complicated the selection of an optimal season for GSV prediction. Thus, a balance between preserving seasonal pattern without sacrificing the temporal resolution needs to be observed. Based on our trade-off analysis, window size = 8 is suitable. With this window size, we proceeded to analyse GSV prediction using different polarizations and estimation methods.

The actual date of the RMSE minimum can be somewhat delayed compared to the date shown on x-axis due to a large window size. A window size of 8 corresponds to approximately three months in the Sentinel-1 time series. The RMSE minimum around December means that the optimal timing for SAR images acquisition is in the winter, around January and February. The peaks around May indicate that the acquisition times in the summer, June or July, are not optimal for GSV prediction in boreal forests when using C-band SAR time series imagery.

#### 4.3.1. Different Regressions over an 8-image Sliding Window

In Figure 9, the window size = 8 is suggested as the sliding window size. Further, to study the influence of different regression methods over sliding windows, we also used MLR and RF in addition to SVR in this subsection. As shown in Figure 10, the seasonal pattern is obvious for all regressions. Seasonal RMSE extrema occurred every summer and winter. The overall RMSE levels and dynamic range were nearly the same for all regression approaches, with MLR underperforming in autumn of 2016. The differences among polarizations were more apparent, favoring VV over VH and overall suggesting combined use of both polarization channels.

**Figure 10:**
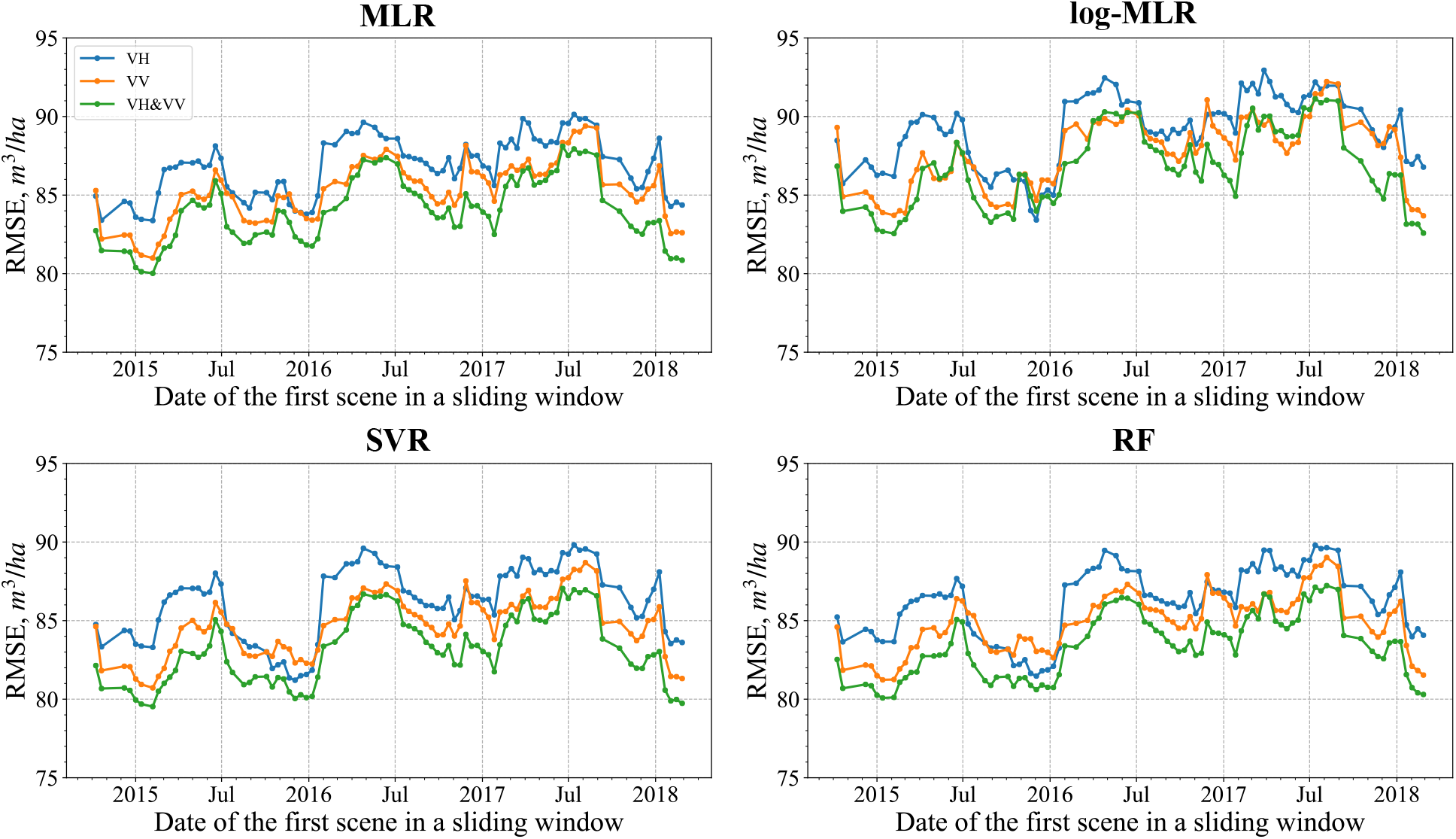
RMSEs of various regression approaches using an 8-image sliding window.

In summary, the sliding window analysis gave a more detailed perspective of the temporal dynamics and facilitates determining the optimal timing for the image acquisition.

## 5. Discussion

### 5.1. On regression approaches

Among the regression approaches SVM, log-MLR and RF showed similar temporal dynamics with no strong differences in GSV prediction. MLR with no log-transformation was less accurate and seems least suitable. RF and SVR produced somewhat greater accuracies than log-MLR in the majority of experiments. MLR with log-transformation of the target variable had somewhat greater variance particularly for larger GSV values, however, it provided the most accurate explanatory model for greater GSV values (Figure 5). No earlier benchmarking studies with C-band data in boreal forests can be found in the literature, while studies with C-band data have been reported for tropical forest (Englhart et al., 2011).

When using only one polarization as the feature variable, no large differences in the accuracies were found, with VV alone giving slightly more accurate GSV predictions than VH alone in non-frozen conditions, and VH being more optimal in frozen conditions. When limiting analysis to stands with only smaller values of GSV, VH-polarization starts to produce consistently more accurate results. Within the small-biomass scenario, VH becomes more sensitive to biomass because cross-pol contains mostly volumetric scattering in forest environment. This discrepancy can be explained by somewhat smaller extinction at VV that contributes to greater sensitivity to large-biomass forest, when the ground is not visible. Under frozen conditions, when the ground is more visible, VV becomes suboptimal compared to VH even when all forest stands are considered, as VV-pol contains both scattering from the ground and volumetric scattering. For both scenarios of GSV values, the use of combined polarizations gave the most accurate predictions. The observation that VH is more suitable is supported by earlier studies, however no analysis for different biomass classes was reported.

### 5.2. The effects of seasonality on GSV prediction

The observation that C-band backscatter images acquired in frozen conditions are more suitable for forest prediction in boreal forests has been reported in earlier studies with ERS by (Pulliainen et al., 1996; Pulliainen et al., 1999) and ASAR (Santoro et al., 2011). Our study confirms this observation on Sentinel-1 images, though the difference was relatively weak when single images were used. Moreover, our feature selection experiments (when 10 most accurate features were selected using Lasso and ik-NN approaches) indicated that adding dry summer images in addition to frozen condition images gives more accurate GSV predictions compared to using exclusively frozen condition images.

Using the sliding window approach suggested in this study for combining features calculated from consecutive Sentinel-1 images as inputs to the regression approach facilitated detection of strong seasonality in GSV predictions compared to using single images. The difference is clearly visible such as in Figure 9. It is important to note that seasonality is not studied here at the level of backscatter (input variable) but rather at the level of GSV prediction (output variable). Using assemblages of features calculated from consecutive images with varying widths of “sliding window” suggests there are optimal timing periods when Sentinel-1 images should be acquired to produce an optimal GSV prediction. Use of a longer “sliding window” permits compensation for the presence of features from suboptimal images and taking advantage of “multitemporal variability” within multi-variable regression models, as features calculated from several consecutive images are used. Our results suggest that even autumn and spring images can be used in principle. This is in contrast with single image retrieval followed by weighting estimates according to accuracy of predictions from individual images (Santoro et al., 2011). In the latter approach, the most accurate strategy is to completely discard autumn and spring images, when the forest biomass C-band SAR to relationship can be most unpredictable. Our results with multivariable regression models indicate that late autumn and early spring datatakes can be also included. For the window size of 8 that corresponds to three months of observations, optimal start can be early as December and as late as February, based on three years of observations.

The use of PCA and several feature selection approaches confirmed that a subset of features calculated using a subset of Sentinel-1 images can be enough to achieve a near optimal GSV prediction. Interestingly, feature selection using mutual information and radiometric contrast led to the least accurate GSV predictions. However, radiometric contrast is the most feasible approach for image selection, particularly when no reference data are available, and is often used for image removal or weighting (Santoro et al., 2011) also Schmullius et al 2012. Lasso was the most accurate option for selecting the features for GSV prediction and approached PCA-based performance in terms of how quickly the near-optimal prediction is achieved. Lasso was closely followed by the ik-NN method. Based on feature ranking with majority voting, Sentinel-1 image #32 acquired on 8 January 2016 was most popular. In addition to images acquired in winter, images acquired in spring and autumn were also sometimes selected. Among the top selected features, VH appeared somewhat more often than VV. No earlier benchmarking studies on feature-selection with C-band data in boreal forests can be found in the literature.

### 5.3. Comparison with previous studies in terms of GSV accuracy and unit-size

Presently, the number of relevant publications on C-band based prediction of GSV is relatively limited compared to L-band and multisensor studies, primarily because of relatively small prediction accuracies. The role of C-band data, particularly Sentinel-1, is quite minor also in the multisensor studies where C-band images are used along with other SAR and also optical datasets (Laurin et al. (2018); Stelmaszczuk-Górska et al. (2018)), although accurate results are reported with a combination of sensors. The results are quite unsatisfactory when using C-band data alone. Our results compare favourably with other published research using C-band SAR data in terms of GSV prediction accuracy (GFOI, 2014; Sinha et al., 2015; Santoro et al., 2011; Kurvonen et al., 1999; Pulliainen et al., 1999;Stelmaszczuk-Górska et al., 2018; Laurin et al., 2018) and are more representative in terms of reference data, feature selection approaches, studied regression models and, compared to the majority of other studies, also use smaller measurement units. Further we compare our experiments with several most representative studies to highlight the relevance of our results.

Use of a parametric semiempirical model in a study by Pulliainen et al. (1999) with set of ERS-1 VV-pol imaged showed that rRMSE of 25–30% can be obtained in boreal forests when the forest block size was larger than 20 ha. Combining ERS-1 images acquired in November and August 1993 was considered optimal which supports our observation that also C-band data acquired in non-winter seasons can be used. Accuracies with smaller sizes of forest stands were much less.

In a study by Kurvonen et al. (1999) with ERS-1 data in boreal forests pursuing GSV prediction using an inversion of a parametric physics-based model, limiting analysis to stands larger than 10 ha led to rRMSE of 34-71%, with the most accurate estimates for mid-December and mid-March. This result is also in agreement with our Sentinel-1 images selected using feature selection approaches. However, incorporating stands with sizes as small as 1 ha increased rRMSE to larger than 120% for separate images. Combining predictions from individual SAR images using multiple regression facilitated a strong reduction in RMSE. However, when ERS-1 images were mixed with JERS (L-band SAR), it was difficult to judge specific contributions, especially because JERS data are much more suitable for GSV prediction.

The BIOMASAR algorithm (Santoro et al., 2011) with the WCM-model inversion at its core used hypertemporal C-band ASAR data at ScanSAR mode (inferior to Sentinel-1 in terms of resolution but exhibiting high radiometric quality due to a large number of looks) for boreal forests GSV prediction with rRMSE between 34.2% and 48.1% at 1 km pixel size. Larger errors were obtained at 100 m, 47.7-60.4% over Swedish test sites, and 70.9-96.2% over Central Siberian test sites.

A study over boreal forests in the Krasnoyarsk region of Russia (Stelmaszczuk-Górska et al., 2018), among other experiments, tested the time series of stripmap data RADARSAT-2 data acquired in regular and ultrafine modes for AGB prediction and achieved rRMSE of 42% and 47% in AGB prediction with a machine learning approach. The mean AGB was approximately 86 t/ha. Time series dynamics were not explored. AGB was predicted using various cross-pol ratios and Haralick features, in addition to average backscatter.

In a recent comprehensive study with several satellite C-band sensors, rRMSE was approximately 90% when predicting AGB (Cartus et al., 2019). However, the error for branch biomass was smaller, 80%. In the case of VV polarization, which included the ERS-2 images acquired in all seasons, the error was greater than 100%. The prediction performance of C-band data depended strongly on the imaging conditions and varied in a large range. The smallest errors were associated with images acquired under stable frozen conditions in winterwhich is similar to our findings. The greatest errors were generally associated with images acquired in periods of snow melt in the spring. This is consistent with our results when using separate SAR images. As ERS-2 has only VV-pol, no separate polarization analysis was possible. It is important to keep in mind that the average and peak values of GSV in Hyytiälä forest were somewhat smaller compared to Reminstorp forest (Cartus et al., 2019).

When comparing our findings to previous studies, as well as to Tomppo et al. (2019), where 30% RMSE was achieved using only 12 Sentinel-1 images, it is important to note that here we used only stand-level backscatter as the independent variable. No ratios or textural features were used, because the aim was to analyse seasonal dynamics and optimal timing of Sentinel-1 data backscatter. It is very likely, that RMSEs would further decrease if other stand-level variables (features) were included in the analysis, in line also with other previous studies ((Tomppo et al., 2019; Wang et al., 2006)).This issue needs extensive additional analysis. Further, we did not pursue the analysis of GSV prediction accuracy as a function of unit area size, but we expect the prediction accuracy will improve as the unit-are size becomes larger, similar to (Pulliainen et al., 1999; Kurvonen et al., 1999; Cartus et al., 2019).

## 6. Conclusions

The utility of very long times series of Sentinel-1 SAR data for boreal forests GSV prediction was tested. We discovered seasonality in GSV prediction with multitemporal C-band time series, suggesting an optimal timing for image collection in early spring and autumn. Several prediction methods gave similar results. The minimum RMSE was 75.6 m^3^/ha (rRMSE 44%) when all 96 images were used. Temporal aggregation of images over one year greatly decreased RMSEs compared to a single image. Feature selection facilitated achieving nearly optimal results using only 10 images (45% relative RMSE) instead of the entire SAR stack, resulting in reduced computational complexity. Among the feature selection approaches, Lasso produced the greatest prediction accuracies. Approaches based on radiometric contrast and mutual information provided much smaller accuracy when the number of selected features was small. When 8-image long “sliding window” time series are used instead of feature selection, relative RMSE of 47% can be achieved if the timing of the image acquisitions is optimal.

Comparison to prior works indicates that no one has studied temporal dynamics of GSV predictions in boreal forest (or elsewhere) in the fashion presented here, using assemblages of consecutive Sentinel-1 images (aka “sliding window” approach). Use of “sliding window” facilitated finding to obtain much stronger annual periodicity in GSV prediction compared to single images, at 1-ha stand-level. Feature selection approaches were used very sporadically, and used methods (correlation, radiometric contrast) were suboptimal as indicated by our experiments. The possibility of using not only winter images, but combining winter and dry-summer images needs further investigation, as our feature selection experiments support the idea of combining corresponding features within multivariate model, where temporal variability is inherently captured.

## Declaring of Competing Interest

None

## Acknowledgements

The Finnish Forest Centre provided the forest resource data of the study. The institutes and universities of the authors provided the computing facilities.

